# Specialized plant biochemistry drives gene clustering in fungi

**DOI:** 10.1101/184242

**Authors:** Emile Gluck-Thaler, Jason C. Slot

## Abstract

The fitness and evolution of both prokaryotes and eukaryotes are affected by the organization of their genomes. In particular, the physical clustering of functionally related genes can facilitate coordinated gene expression and can prevent the breakup of co-adapted alleles in recombining populations. While clustering may thus result from selection for phenotype optimization and persistence, the extent to which eukaryotic gene organization in particular is driven by specific environmental selection pressures has rarely been systematically explored. Here, we investigated the genetic architecture of fungal genes involved in the degradation of phenylpropanoids, a class of plant-produced secondary metabolites that mediate many ecological interactions between plants and fungi. Using a novel gene cluster detection method, we identified over one thousand gene clusters, as well as many conserved combinations of clusters, in a phylogenetically and ecologically diverse set of fungal genomes. We demonstrate that congruence in gene organization over small spatial scales in fungal genomes is often associated with similarities in ecological lifestyle. Additionally, we find that while clusters are often structured as independent modules with little overlap in content, certain gene families merge multiple modules in a common network, suggesting they are important components of phenylpropanoid degradation strategies. Together, our results suggest that phenylpropanoids have repeatedly selected for gene clustering in fungi, and highlight the interplay between gene organization and ecological evolution in this ancient eukaryotic lineage.

## Introduction

Genome organization is intimately linked to the trajectory of organismal evolution. The impacts of genome organization on prokaryotic evolution in particular have been extensively studied (Baquero 2004), and increasingly, the organization of genes in eukaryotic genomes is also recognized to affect organismal fitness and evolution. For example, the spatial clustering of functionally related genes may enable the compartmentalization and optimization of phenotypes through coordinated gene expression (Hurst et al. 2002; Al-Shahrour et al. 2010; McGary et al. 2013). Similarly, the formation of loci composed of co-adapted alleles can facilitate the inheritance of locally adapted ecotypes within recombining populations over short time periods (Yeaman 2013; Holliday et al. 2016). Rather than resulting from non-adaptive processes, such as genetic hitchhiking, the persistence of such organizational patterns in eukaryotic genomes suggests they may instead result from natural selection (Hurst et al. 2002; Lynch 2007). However, the extent to which eukaryotic genome organization is driven by specific environmental selection pressures, especially over macroevolutionary timescales, is not clear.

Organized genome structure is particularly apparent in fungi, a lineage of eukaryotic microorganisms that perform critical ecosystem services and biomass transformation, and also impact plant and animal health (Peay et al. 2016). Fungal genomes are more or less replete with metabolic gene clusters (MGCs) composed of genes encoding enzymes, transporters and regulators that participate in specialized metabolic processes such as nutrient acquisition, competition and defense (Wisecaver et al. 2014). Although MGCs are far more rare in eukaryotes compared with bacteria, fungal MGCs exhibit similarly sparse and disjunct phylogenetic distributions among distantly related species with overlapping niches (Greene et al. 2014; Dhillon et al. 2015; Glenn et al. 2016). This ecological pattern of distribution suggests that conserved combinations of genes may be signatures of ecological selection in fungal genomes.

Fungal MGCs encoding the production of specialized, or secondary, metabolites (SMs) have been studied extensively (Hoffmeister and Keller 2007) and more recently, several reports suggest that adaptations to degrade plant SMs are also encoded in MGCs (Shanmugam et al. 2010; Greene et al. 2014; Wang et al. 2014; Kettle et al. 2015; Glenn et al. 2016). Plant SMs mediate important biotic and abiotic interactions, including the exclusion of fungal pathogens, the establishment of mutualisms, and the rates of nutrient cycling long after the plant has died. The largest group of plant SMs are the phenylpropanoids, which not only contribute to constitutive and inducible chemical defenses, but are also the main barriers to wood decay in terrestrial ecosystems (Floudas et al. 2012), and costly inhibitors of lignocellulose biofuel production (Jönsson and Martín 2016). As the primary colonizers of plant material in natural and artificial environments, fungi are frequently in contact with phenylpropanoids, and must often mitigate their inhibitory effects in order to grow. Common fungal adaptations to phenylpropanoid toxicity include detoxification through sequestration, excretion and degradation (Mäkelä et al. 2015). Despite the characterization of many phenylpropanoid degradation pathways in fungi, the genomic bases of these pathways are largely unknown (Mäkelä et al. 2015), precluding the use of currently available algorithms to investigate whether or not these metabolic processes are encoded in MGCs (Wisecaver et al. 2014; Weber et al. 2015).

Here, we developed a novel algorithm based on empirically derived models of fungal genome evolution in order to systematically identify clusters of genes putatively involved in phenylpropanoid degradation, enabling us to test the hypothesis that selection pressures from plant SMs impact genome organization across disparate fungal lineages. Using a database of 556 fungal genomes representing 481 species, we detected 1168 MGCs and many conserved combinations of MGCs that putatively degrade a broad array of phenylpropanoids. We then tested for associations between MGCs and various fungal ecological lifestyles, and found that the presence of certain MGCs was enriched in plant pathotrophs and saprotrophs. While many clusters appear to have evolved independently, we identified several gene families that are commonly associated with diverse MGCs, suggesting they play important roles in phenylpropanoid catabolism. These results suggest that phenylpropanoids are drivers of gene organization in plant-associated fungi, and that MGCs may in turn determine patterns of fungal community assembly on both living and decaying plant tissues.

## Results

### Diverse candidate gene clusters are associated with phenylpropanoid degradation

Using 27 different “anchor” gene families involved in phenylpropanoid degradation as separate queries (Methods; Supplementary Table 1), we searched 556 genomes from 481 fungal species for clusters associated with each anchor gene family (Methods; Supplementary Figure 1, Supplementary Table 2). After removing those clusters containing genes known to exclusively participate in fungal secondary metabolite biosynthesis (Supplementary Table 3; Supplementary Table 4), as well as gene clusters with fewer than 4 genes, we found evidence of unexpected clustering in 13 anchor gene families, which we defined as separate cluster classes. We identified a total of 1168 clusters distributed across 363 fungal genomes (Figure 1, Supplementary Figure 2, Supplementary Table 5). 31 of these clusters belonged to multiple clusters classes (i.e., contained multiple anchor genes) (Supplementary Table 3). When accounting for multiple occurrences of homologous clusters (as defined by cluster model; Methods) among individuals of the same species, this total figure was reduced to 962 clusters distributed across 309 fungal species. The number of cluster models per cluster class varies from 1 to 8, and clusters assigned to different models generally do not overlap in terms of content (Supplementary Table 6). Additionally, clusters typically contained very few intervening genes (Supplementary Figure 3). Clusters are disproportionately distributed across fungal lineages: for example, Pezizomycotina comprises 50.3% of the species in the analysis yet contains 89.4% of all clusters, while Agaricomycetes represent 22.9% of species, but have only 4.7% of clusters (Supplementary Table 7). However, homologous clusters assigned to the same model are typically found distributed across different taxonomic classes (Supplementary Table 7).

**Figure 1:**
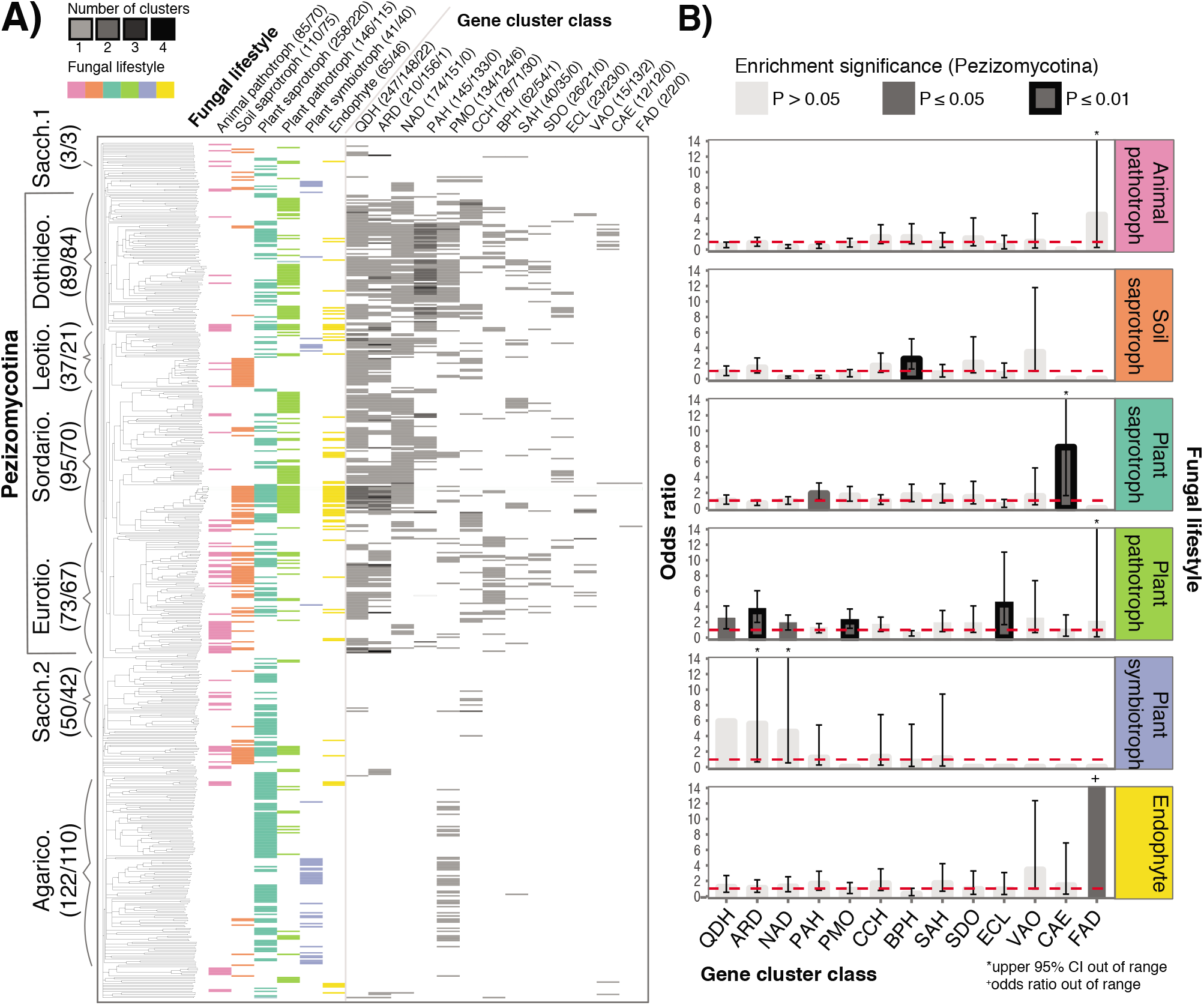
Associations between gene cluster distributions and fungal lifestyle. a) A phylogeny of 556 fungi (representing 481 species) based on pairwise microsyntenic similarity is shown to the left, annotated by taxonomic class. All taxonomic classes have been abbreviated by removing “-mycetes” suffixes. A matrix indicating the ecological lifestyle (s) associated with each fungus is shown to the immediate right of the phylogeny, followed by a heatmap indicating the number of putative phenylpropanoid-degrading gene clusters in each genome, for each cluster class. Numbers in brackets following the taxonomic class and ecological lifestyle headers correspond to the number of genomes and species within those categories. Numbers in brackets following cluster class headers indicate the total number of clusters assigned to that class, the number of species with at least 1 cluster, and the number of clusters that overlap with clusters from another cluster class. b) Odds ratios representing the strength of the association between cluster presence and fungal ecological lifestyle are shown for each of 13 gene cluster classes and 6 lifestyles, using data at the species level from the Pezizomycotina. Dotted red lines indicate an odds ratio of 1. Dark grey bars indicate enrichment below a significance level of 0.05, while black outlines indicate enrichment below a significance level of 0.01. Error bars indicate the 95% confidence interval (CI) for each odds ratio measurement. CIs of 0 are not shown. The color-coding of ecological lifestyle is consistent across the entire figure.

In order to benchmark our approach, we searched for previously characterized clusters containing quinate 5-dehydrogenase (QDH) and stilbene dioxygenase (SDO) homologs. All 8 previously predicted clusters from the SDO cluster class (Greene et al. 2014) and all 31 described clusters from the QDH class (Hane et al. 2007; Shalaby et al. 2012) were recovered. Additionally, while this study was being prepared, a single cluster from the pterocarpan hydroxylase (PAH) class was identified elsewhere (Pigné et al. 2017). To our knowledge, the remaining 1128 clusters identified in this analysis are described here for the first time. We expect some false negative error in our detection but not significant false positive to arise due to variation in the quality of genome assemblies and annotations. Some false positive error in our detection may, however, arise from the fact that beyond the function of the anchor gene, we did not require any genes to be demonstrably involved in phenylpropanoid degradation because the loci involved in such pathways are for the most part unknown (Mäkelä et al. 2015). Although we removed clusters with genes known to exclusively participate in biosynthetic secondary metabolism (Supplementary Table 4), it is still possible that some of the remaining clusters participate in biosynthetic reactions, as it is often difficult to definitively distinguish between biosynthetic and catabolic processes.

### Clustered homolog groups, including those distributed across cluster classes, are enriched for primary and secondary metabolic functions

Despite the lack of functional constraints on cluster composition, clustered homolog groups were, enriched for 7 KOG processes primarily related to either primary or secondary metabolism, including “Energy production and conversion” and “Secondary metabolite biosynthesis, transport and catabolism”, the two largest and most significantly enriched categories (Supplementary Table 8). The KOG process “Transcription” was the only non-metabolic process enriched in clustered homolog groups. Homolog groups present in more than one cluster class are also enriched for multiple functional categories related to primary and secondary metabolism (Supplementary Table 9). Conserved domains from the Pfam database that are present among shared homolog groups include transport and transcription-related domains, cytochrome P450 domains, and domains related to functional group modification (e.g., dehydrogenase, transferase, amidase and decarboxylase; Supplementary Table 10).

### Phenylpropanoid degradation gene clusters are enriched in Pezizomycotina species with plant-associated lifestyles

We limited our exploratory enrichment analyses to the Pezizomycotina, as they contained the vast majority of identified clusters. Clusters from 5 cluster classes are enriched in plant pathotrophs, including those containing the anchor genes naringenin 3-dioxygenase (NAD) and epicatechin laccase (ECL) that are know to participate in flavonoid degradation (Figure 1). An additional 3 cluster classes are enriched in other plant-associated lifestyles, such as plant saprotrophs and endophytes. The benzoate 4-monooxygenase (BPH) cluster class is enriched in soil saprotrophs. No cluster classes are enriched in plant symbiotrophs or animal pathotrophs. Similar patterns of lifestyle-dependent enrichment are observed when comparing Pezizomycotina species with specific cluster models to those without any cluster models (Supplementary Figure 4). Notably, at least one model in each of 12 cluster classes is enriched in a plant-associated lifestyle. Aromatic ring dioxygenase (ARD) models are most often enriched in plant-associated lifestyles: model 1 is enriched in plant symbiotrophs and plant pathotrophs; model 2 is enriched in plant pathotrophs, and model 5 is enriched in endophytes. Although ecological enrichments highlight predominant trends, clusters across all anchor genes are typically associated with diverse lifestyles, in part because any given fungal species can be associated with multiple ecological lifestyles.

We observed that 207 fungal species possessed more than one cluster model. These fungi were assigned to multi cluster model profiles (MCMPs) based on the combinations of cluster models found in their genomes (Methods). 139 fungal species were distributed across 16 distinct MCMPs with 5 or more species (Figure 2). Fungi from the same MCMP tend to be closely related; however, 13 MCMPs contain fungi from different taxonomic orders, and 6 of these contain fungi from different taxonomic classes. We also limited this exploratory enrichment analysis to Pezizomycotina species and found that 4 multi-cluster profiles are enriched in species with plant-associated lifestyles, 1 is enriched in animal pathotrophs and endophytes, and 1 is enriched in soil saprotrophs (Figure 2).

**Figure 2:**
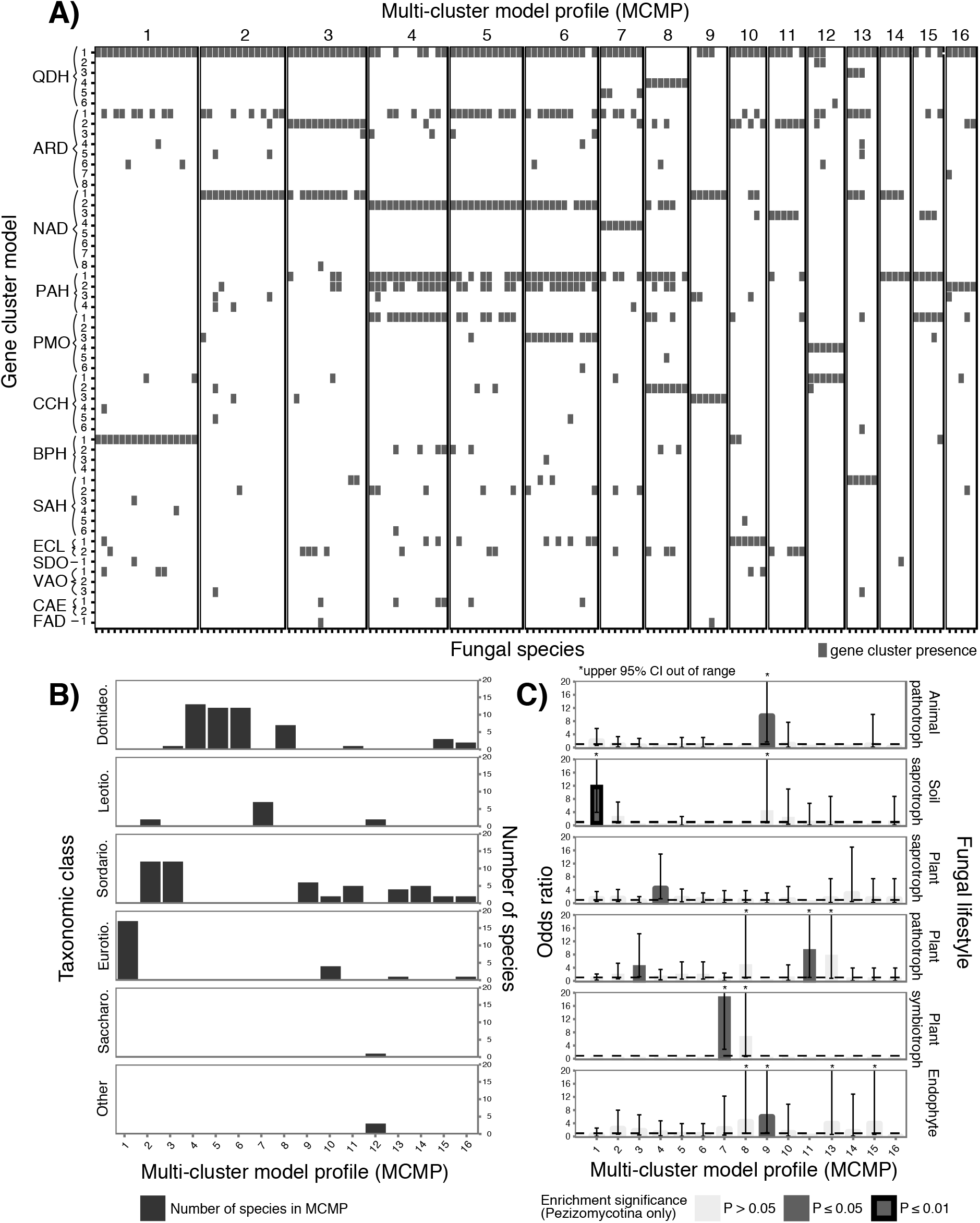
Combinations of putative phenylpropanoid-degrading gene clusters in fungal genomes. a) A matrix describes the presence/absence of various homologous clusters (as determined by cluster model; rows) in the genomes of 139 fungal species (columns). Species are grouped into 16 multi-cluster model profiles (MCMPs) based on similarities in the combinations of clusters found in their genomes. b) A bar chart depicts the number of fungal species from each MCMP per taxonomic class. c) Odds ratios representing the strength of the association between MCMP and fungal ecological lifestyle are shown, using data at the species level from the Pezizomycotina. Dotted black lines indicate an odds ratio of 1. Dark grey bars indicate enrichment below a significance level of 0.05, while black outlines indicate enrichment below a significance level of 0.01. Error bars indicate the 95% confidence interval (CI) for each odds ratio measurement. CIs of 0 are not shown. Enrichment data is not shown for MCMP 12, as fewer than 5 fungi from the Pezizomycotina are assigned to this MCMP.

### Homolog groups are found in modules of co-clustered genes

We visualized structural associations among all clustered homolog groups as a network where nodes represent homolog groups, and edges represent the co-occurrence of homolog groups in a cluster (Figure 3). A number of homolog groups are found in clusters belonging to different cluster classes (39 in total; Supplementary Table 10), which resulted in a common network for all cluster classes. Shared homolog groups are typically among the most highly connected nodes in this network (i.e., they co-occur frequently with a diverse set of homolog groups; Supplementary Table 10). The modularity of the network is significantly greater than that observed in networks of similar size with randomly distributed edges (Supplementary Table 11). This high degree of modularity reflects a large number of homolog groups that are unique to each cluster class (238 in total). Some homolog groups, while not shared across cluster class, are highly connected because they are present in different cluster models (e.g., see homolog group 1001 in Figure 4).

**Figure 3:**
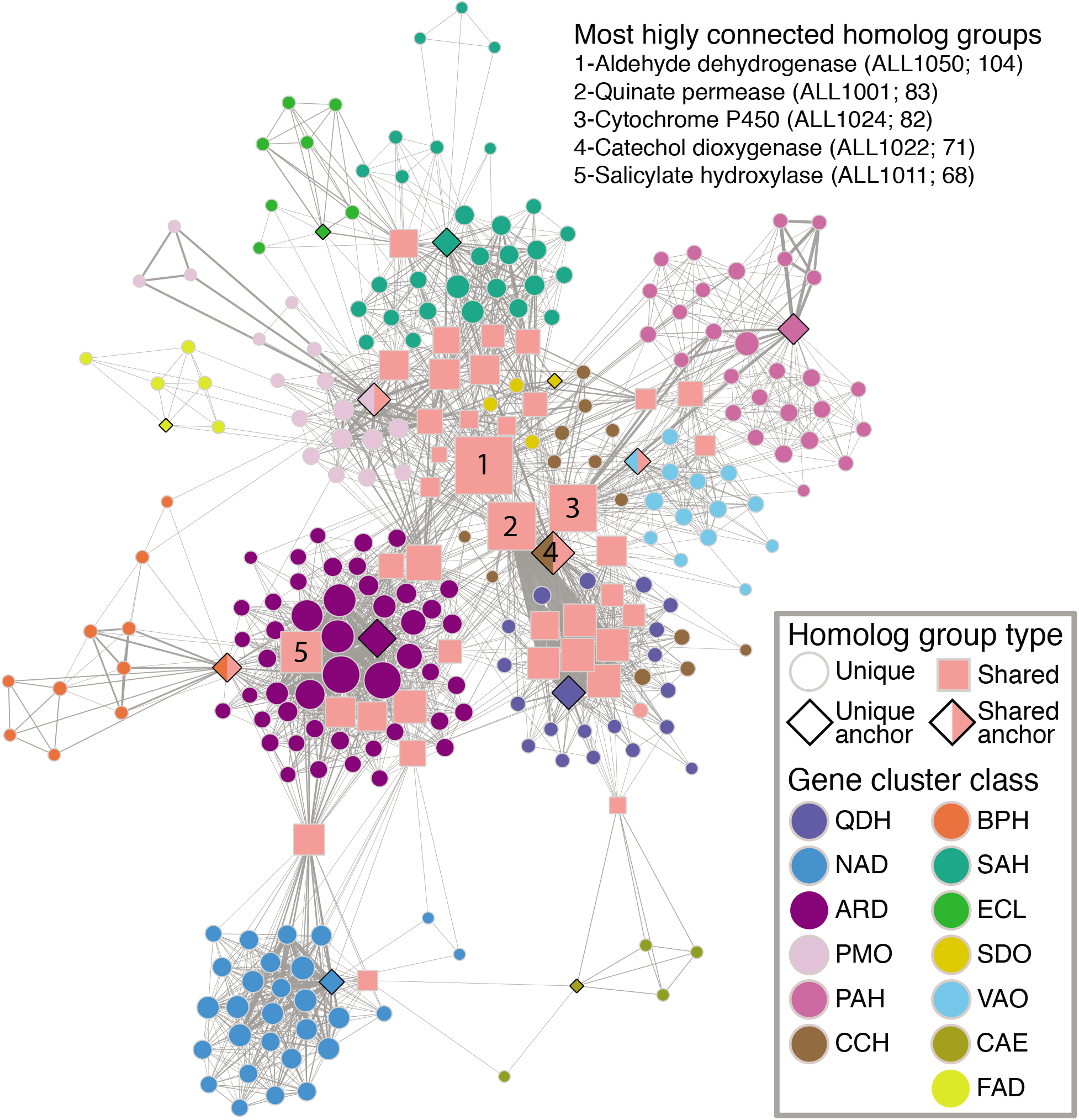
Co-occurrence network of homolog groups in putative phenylpropanoid-degrading gene clusters. Each node represents a homolog group found in a putative phenylpropanoid-degrading gene cluster. All nodes are color-coded by the cluster class in which they are found, except for those homolog groups found associated with multiple cluster classes (i.e., shared), colored pink and shaped as squares. Nodes representing anchor gene families used to retrieve the clusters are shaped as diamonds, and those anchor gene families that are shared across multiple cluster classes are additionally colored pink. Edges symbolize the co-occurrence of two homolog groups in the same gene cluster, while edge width is proportional to the frequency of that occurrence. Node size is proportional to the number of connections emanating from that node. The proximity of nodes to one another is proportional to the number of shared connections. The annotations, followed by the code names and number of unique connections in parentheses, of nodes with the greatest number of connections (i.e., associated with the greatest diversity of homolog groups) are indicated in the top right-hand corner of the network.

**Figure 4:**
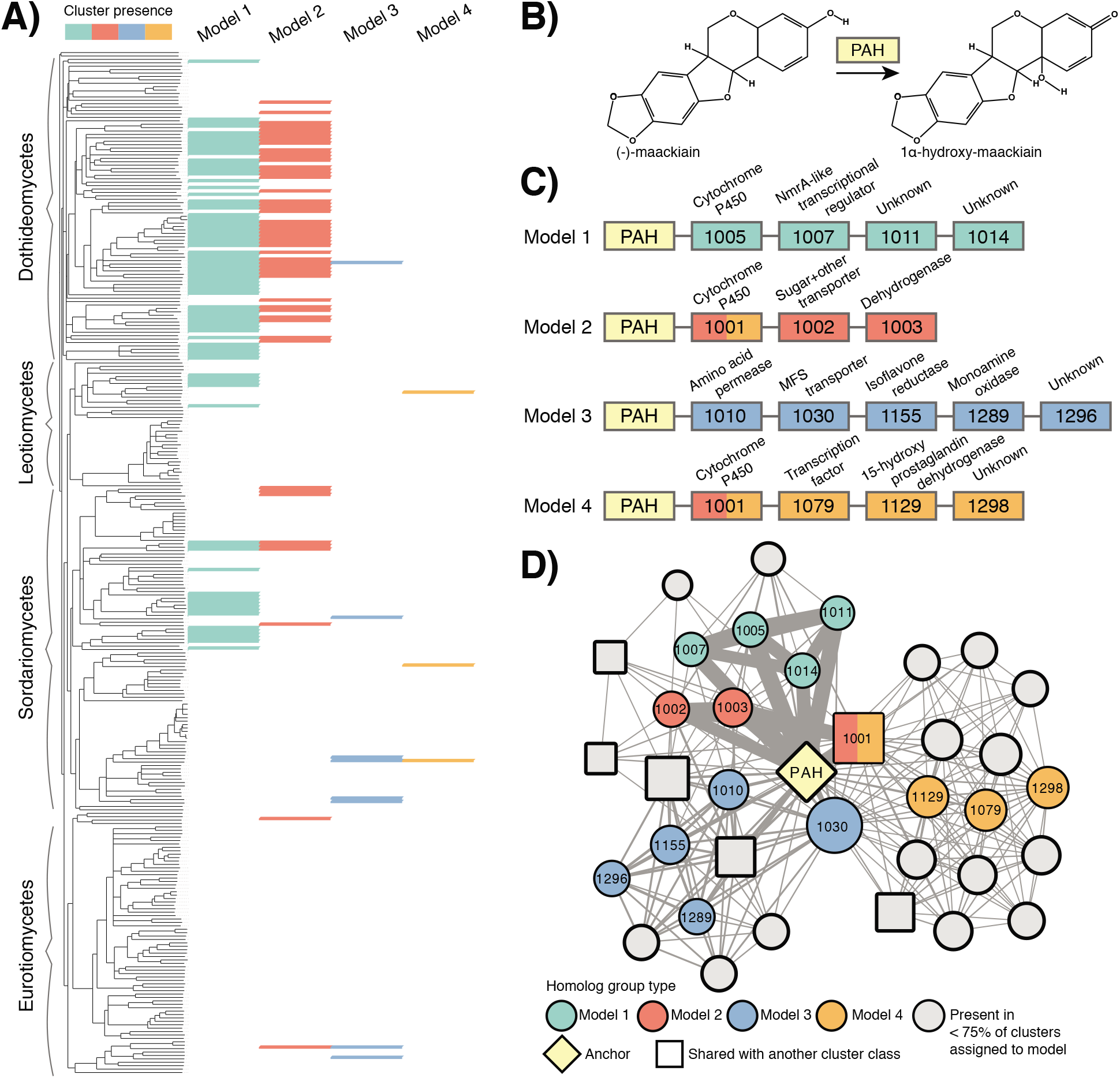
The distribution of pterocarpan hydroxylase (PAH) clusters in fungi. a) A phylogeny of the Pezizomycotina based on pairwise microsyntenic distance is shown to the left, annotated by taxonomic class. The presence/absence of clusters assigned to the 4 PAH cluster models is indicated in the matrix to the right of the phylogeny, color-coded by cluster model. b) A simplified schematic of one of several reactions catalyzed by PAH. c) Homolog groups present in ≥75% of clusters assigned to a given cluster model are depicted as boxes color-coded by cluster model, while PAH homologs are indicated in yellow. The four-digit code and predicted annotation are indicated for each homolog group. D) The depicted network follows the conventions specified in Figure 2. Homolog groups present in ≥75% of clusters assigned to a given cluster model are color-coded by cluster model and inscribed with their code, while others are colored gray. Nodes representing homolog groups present in clusters from other cluster classes are drawn as squares.

### Results spotlight (could be displayed as a Box, combining results and discussion): Pterocarpan hydroxylase clusters may encode uncharacterized strategies for flavonoid degradation

The production of toxic SMs by plants is an ancient and effective strategy for deterring fungal growth. Isoflavonoids, for example, are a critical component of legume (Fabaceae) defenses against fungi. In response, some legume pathogens have evolved degradative metabolic pathways to detoxify isoflavonoids, thus facilitating host colonization (Enkerli et al. 1998). Pterocarpan hydroxylase (PAH) is involved in the degradation of pterocarpan and medicarpin isoflavonoids, and can contribute to virulence on susceptible plant hosts (Enkerli et al. 1998). We found 133 clusters in the PAH class, distributed across 94 fungal species with diverse ecological lifestyles (Figure 4). Homology among these clusters is best described by 4 models containing many of the canonical functions found in fungal MGCs. Models 1, 2 and 4 all contain cytochrome P450s, a large family of enzymes involved in detoxification mechanisms in fungi and other organisms (Lah et al. 2011), while models 2 and 3 contain at least one transporter and model 4 contains a transcription factor. Model 3 contains a homolog of isoflavone reductase, an enzyme used in isoflavonoid biosynthesis in plants, but may also be involved in isoflavonoid detoxification pathways (Höhl et al. 1989). The widespread distribution of PAH clusters suggests they are involved in the degradation of flavonoids found outside the Fabaceae, and the enrichment of model 1-type clusters in plant saprotrophs suggests that flavonoids that persist in plant tissues may impact fitness in saprophytic environments. However, the genetics underlying isoflavonoid biosynthesis and degradation, and flavonoid metabolism in general, are not well understood for fungi, despite the ecological importance of this chemical family.

A PAH homolog in *Alternaria brassisicola* found in a cluster assigned to model 1 was recently shown to contribute to the accumulation of DHN melanin in cell walls, possibly through catalyzing melanin polymerization or the cross-linking of melanin to fungal cell walls (Pigné et al. 2017). Melanins are a class of complex phenolic polymers that can serve to quench oxidizing radicals, and play a role in fungal pathogenicity and survival in harsh environments (Bayry et al. 2014). Intriguingly, plant flavonoids can be polymerized in planta and in vitro into melanin by fungal laccases (Fowler et al. 2011) and possibly by monooxygenases similar to PAH (Desentis-Mendoza et al. 2006). As the repurposing of host metabolites as substrates for the production of SMs is not unprecedented in fungi (Schmaler-Ripcke et al. 2009), we suggest that this PAH cluster in particular is a good candidate for exploring hypotheses of cross talk between specialized metabolic pathways.

## Discussion

### Candidate phenlypropanoid degradation MGCs found at over 1000 loci with unexpectedly conserved synteny

Fungi are the most ecologically significant decomposers of plant biomass, and represent some of the world's most devastating plant pathogens (Peay et al. 2016). The presence of catabolic genes targeting plant SMs in fungal genomes has often been linked to species ecology and the ability to colonize different types of plant tissues (Floudas et al. 2012). Much less is known about how these genes are organized in fungal genomes, and how their organization impacts or may be affected by fungal evolution and ecology. The genomes of many fungi tend to undergo extensive and frequent rearrangements (Hane et al. 2011; Plissonneau et al. 2016). By contrast, MGCs we detected are often simultaneously present in lineages that diverged hundreds of millions of years ago (Floudas et al. 2012) (Figure 1). As combinations of genes that persist over long periods of evolutionary time are likely to be maintained by natural selection (Hurst et al. 2002; Baquero 2004; Muto et al. 2013) and often encode proteins that are functionally related (Szklarczyk et al. 2014; Zhao et al. 2014), we hypothesize that the conserved combinations of phenylpropanoid degrading genes detected here encode adaptive metabolic phenotypes and are signatures of selection on gene organization.

The most well studied phenotype conferred by clustering is the improved coordination of gene transcription (Hurst et al. 2002). Clustering facilitates gene transcription by allowing co-regulation through local chromatin modifications (Shwab et al. 2007), the sharing of promoters (Davila Lopez et al. 2010), and the avoidance of topological constraints on DNA encountered during transcription (Tsochatzidou et al. 2016). In addition to synchronising cellular responses, coordinated gene transcription may also decrease the accumulation of toxic intermediate metabolites, such as those produced during phenylpropanoid degradation (Greene et al. 2014; Mäkelä et al. 2015), by optimizing enzyme stoichiometry and improving metabolic flux (McGary et al. 2013). Gene clustering may additionally be selected for because it increases the capacity for recombining populations to evolve. When genes contributing to the same trait are clustered, selection for that trait is more efficient because of decreased selective interference between genes at that locus (Pepper 2003). Clustering can also prevent the breakup of co-adapted alleles in the face of gene flow (Yeaman 2013), which may enable local metabolic adaptation of fungal populations distributed across cryptic niches.

Similarly, clustering increases the probability of propagating adaptive combinations of genes through both vertical (Pepper 2003) and horizontal (Lawrence and Roth 1996; Baquero 2004) transfer. Many of the clusters observed in this study have discontinuous phylogenetic distributions that are consistent with evolutionary scenarios such as vertical inheritance coupled with extensive loss, convergence, duplication, or horizontal gene transfer (HGT). HGT of MGCs is predicted to be associated with rapid adaptive changes to phenotypes, including increased capacity for host colonization (Dhillon et al. 2015; Glenn et al. 2016) and nutrient acquisition (Slot and Hibbett 2007). For example, MGCs in bacteria are frequently dispersed by HGT among species inhabiting the same environment or host (Baquero 2004). HGT is positively influenced by relatedness in bacteria (Andam and Gogarten 2011) and fungi (Wisecaver et al. 2014; Gluck-Thaler and Slot 2015), as well as by shared ecological niche (Smillie et al. 2011). Given that the distributions we observed here may be at least partly HGT-driven (Greene et al. 2014), the role of HGT in the dispersal of putative phenylpropanoid degrading MGCs must be explicitly tested with phylogenetic methods (Gluck-Thaler and Slot 2015).

We detected the vast majority of clusters in sub-phylum Pezizomycotina. Among fungi, the Pezizomycotina are known to possess highly clustered genomes, while phylum Basidiomycota and sub-phylum Saccharomycotina have much fewer (Wisecaver et al. 2014). Such differences may be due modes of chromosomal evolution unique to the Pezizomycotina that involve extensive rearrangements that could facilitate cluster formation (Hane et al. 2011). The large population sizes attributed to many Pezizomycotina lineages could then serve to increase selection efficiency for clusters (Lynch and Walsh 2007) such that they would be maintained. Fungi in the Pezizomycotina are also thought to experience more HGT than those in other phyla (Wisecaver et al. 2014), which may facilitate cluster dispersal.

### Gene cluster distributions suggest ecological adaptation

While certain clusters have distributions suggestive of lineage-specific bias (Supplementary Table 7), these conserved combinations of genes were ultimately detected by their unexpected phylogenetic distributions that conflicted with lineage-specific phylogenetic signal, indicating that phylogeny is not the sole determinant of cluster distributions. Indeed, gene organization is often an important determinant of fitness and may be differentially selected across ecological niches (Kirkpatrick and Barton 2006; Yeaman 2013). Our analysis suggests that the presence of putative phenylpropanoid degrading MGCs in the Pezizomycotina is associated with ecological lifestyle (Figure 1), although this does not of course preclude other traits, such as host preference, that might also affect cluster distributions. Intriguingly, two of the five cluster classes enriched in plant pathotrophs (NAD and ECL) are predicted to target flavonoids, which are important components of plant defense against fungal pathogens. Such clusters may contribute to pathogen fitness, as recent reports suggest that the degradation of host defense compounds contributes quantitatively to plant pathogen virulence and reproduction (Hammerbacher et al. 2013; Kettle et al. 2015). Other cluster classes enriched in plant pathotrophs, including QDH, ARD and phenol 2-monooxygenase (PMO), are predicted to target less derived molecules, and may participate in the downstream degradation of more complex phenylpropanoids, such as lignin (Jönsson and Martín 2016). Catabolic clusters may also confer fitness benefits to saprotrophs living in plant tissues or soil by enabling the degradation of phenylpropanoids that inhibit fungal colonization of organic matter (Floudas et al. 2012). Indeed, many of the cluster classes enriched in saprotrophs, including ferulic acid esterase (CAE) and BPH are predicted to target phenylpropanoids commonly found in decaying plant material and soils (Mäkelä et al. 2015). We also found that individual cluster models are enriched in plant-associated lifestyles, giving insight into the specific patterns of gene organization favoured by natural selection (Supplementary Figure 4). Clusters from different models were rarely enriched in more than one ecological lifestyle, possibly reflecting differential selection pressures imposed by phenylpropanoids encountered in different ecological contexts.

### Cluster model co-occurrences may reflect simultaneous selection by multiple plant metabolites

The modularity of the structural network of clustered homolog groups suggests that different cluster classes carry out semi-independent functions (Wagner 1996), and may reflect differences in phytochemicals processed by each cluster class (Figure 3). Minimally, the structuring of clusters as independent modules with little overlap in content indicates that cluster classes have largely evolved independently of one another, and that selection for gene organization is likely to have occurred multiple times.

We sorted different fungal species into multi-cluster model profiles (MCMPs) based on the combinations of clusters found in their genomes (Figure 2), raising the intriguing possibility that conserved cluster repertoires may be important for determining ecological phenotypes, similar to how combinations of pathogenicity islands in bacteria determine host range (Bouyioukos et al. 2016). While certain MCMPs were only present in specific fungal classes, others were present in fungi from different taxonomic classes and orders, and certain MCMPs are enriched in Pezizomycotina species with specific plant-associated lifestyles (Figure 2). Plants typically do not produce a single chemical in isolation; rather, defense metabolites are released as mixtures, either with metabolites from different pathways, or with metabolic precursors from the same pathway, or both (Gershenzon et al. 2012). MCMPs may reflect the compositions of SM mixtures encountered during the colonization of specific hosts, or possibly temporal differences in defense compound pressure (Hammerbacher et al. 2013), and may underlie assembly patterns of fungal communities on different hosts and substrates. A detailed investigation of the full complement of degradative MGCs from fungi colonizing the same plant host would be ideal for testing this hypothesis.

### Do gene clusters bridge plant and fungal metabolisms?

In order to infer which clustered homolog groups may play outsized roles in fungal strategies for phenylpropanoid degradation, we ranked each group by the number of non-redundant co-occurrences they have with other clustered homolog groups, and identified homolog groups present in different cluster classes (Figure 3; Supplementary Table 10). We predict that the homolog groups with the greatest number of co-occurrences (i.e., the most highly connected, Figure 3), as well as those found associated with the greatest number of cluster classes, have been repeatedly recruited to diverse catabolic processes and may encode key strategies for phenylpropanoid degradation in fungi. Notably, some of the shared homolog groups, like those with members encoding cytochrome P450 enzymes (see groups ALL1024, ALL1032, ALL1115 in Supplementary Table 10) and multidrug transporters (ALL1122, ALL1127), are known to be associated with evolutionary adaptability and often underlie fungal detoxification strategies of diverse chemicals (Lah et al. 2011; Mäkelä et al. 2015).

Fungi often exclusively acquire carbon from plant-derived substrates. Intriguingly, the cellular processes associated with shared homolog groups also suggest selection to integrate phenylpropanoid degradation products into central fungal metabolism. Other shared homolog groups have members normally associated with core fungal metabolism, for example, carbohydrate transporters (MFS monocarboxylate transporter: ALL1065; Sugar (and other) transporter: ALL1071, ALL1103), enzymes that produce substrates for aromatic amino acid biosynthesis (3-dehydroshikimate dehydratase: ALL1002), and enzymes that participate in aromatic amino acid degradation (fumarylacetoacetate hydrolase: ALL1055). Metabolic processes, such as those often encoded in clusters (Muto et al. 2013; Greene et al. 2014), are predicted to evolve through the recruitment and subsequent specialization of duplicated enzyme-encoding genes from existing metabolic pathways (Jensen 1976). Shared homolog groups thus offer evidence that genes involved in SM degradation as well as those involved in core fungal metabolism have been recruited from existing pathways to MGCs, as has been suggested elsewhere (Greene et al. 2014). Combinations of genes in MGCs may ultimately serve to connect diversifying chemical environments to a stable metabolic network, acting as a bridge between specialized plant metabolism and core fungal metabolism, and enabling fungal growth under what would otherwise be challenging conditions.

The prevalence of putative phenylpropanoid-degrading MGCs in plant-associated fungi suggests that specialized plant metabolism is a strong source of selection on fungal gene organization. Based on their distribution, we hypothesize that the MGCs detected here encode selectable phenotypes that promote the colonization of plant substrates by fungi. Further functional characterization of these loci will serve to accelerate discovery of metabolic processes that are of great interest not only for lignocellulosic biofuel production, but also for our understanding of the evolutionary dynamics between gene organization and ecological adaptation.

## Materials and methods

### Data acquisition, annotation and software specifications

Publically available data from 556 assembled fungal genomes and predicted proteomes were downloaded from various sources (Supplementary Table 2). While data from all genomes was used to generate the presented results, specific results from the 209 unpublished genomes have been redacted, and assigned generic names based on taxonomic class. For each gene with alternative transcripts in the set of genomes, the predicted amino acid sequence associated with the longest of such transcripts was used for cluster detection, and the rest were discarded. Ecological metadata of all 556 genomes were compiled from various sources (Supplementary Table 2), including the community-curated FUNGuild database (last accessed 09/09/16) (Nguyen et al. 2016) and the U.S. National Fungus Collection database (last accessed April 1st, 2017) (Farr and Rossman 2017). The naming of ecological lifestyles follow the framework established by FUNGuild (Nguyen et al. 2016).

Amino acid sequence searches with cutoffs of 30% identity, 50 bitscore and where the length of the target sequence was 50-150% of the query sequence, were performed with using USEARCH v8.0.1517's UBLAST algorithm (additional parameters: evalue cutoff = 1e-5, accel = 0.8) (Edgar 2010) or with BLASTp v2.2.25+ (additional parameters: evalue cutoff = 1e-4) (Altschul et al. 1990). Homology was determined using OrthoMCL v2 with an inflation value of 1.5 (Fischer et al. 2011). All amino acid sequences from clustered homolog groups were assigned to orthologous groups from the fuNOG database (last accessed 03/18/16) (Huerta-Cepas et al. 2015) using HMMER3 (Eddy 2011). Clustered sequences were additionally annotated by PFAM and GO term using InterProScan 5 v5.20-59.0 (Jones et al. 2014). Predicted functional annotations and KOG processes for each clustered homolog group were based on the most frequent fuNOG annotation assigned to proteins within that group. All phylogenetic trees were visualized using ETE v3 (Huerta-Cepas et al. 2016) and all other graphs were visualized using the ggplot2 package in R (Wickham 2016). All data used to generate the presented figures can be found in Supplementary Table 12.

### Construction of the microsynteny tree

The evolution of microsynteny (gene content conservation over small genomic distances) does not necessarily recapitulate phylogenetic relationships determined by models of sequence evolution. In order to assess the unexpectedness of cluster distributions within a phylogenetic framework based on microsynteny conservation, we used an approach similar to Snel et al. (1999) (Snel et al. 1999) to construct a species tree based on microsyntenic distance, or pairwise comparisons of gene content conservation (Supplementary Figure 1). Distances based on microsynteny are necessarily bounded by some maximum value that is often rapidly reached as microsynteny degrades, and thus the ability to resolve phylogenetic relationships decreases as evolutionary distance increases (Rogozin et al. 2004). Nevertheless, the species relationships that we recovered in general correspond well with those obtained using conventional phylogenetic methods (Schoch et al. 2009; Spatafora et al. 2016).

To construct the microsynteny tree, up to 10 genes upstream and downstream of all homologs from a randomly selected gene family were retrieved (designated “gene neighborhood”), then had their amino acid sequences compared against one another using UBLAST, and subsequently were sorted into homolog groups using OrthoMCL. For each pairwise neighborhood comparison, the total number of shared orthologs was divided by the size of the smallest neighborhood, subtracted from 1, and designated as pairwise syntenic distance. Pairwise syntenic distances thus range from 0 (all homolog groups are shared within a neighborhood) to ~0.95 (only one gene, the randomly selected query, is shared). 1000 neighborhoods were randomly sampled in this way. A neighbor-joining tree was constructed from the distance matrix of the median pairwise syntenic distance for each pairwise genome comparison using the ape package in R (Paradis et al. 2004). The original dataset was sampled with replacement to obtain 100 bootstrap pseudo-replicate trees that were then used to calculate bootstrap support on the original tree using the –z switch in RAxML v8.2.0 (Stamatakis 2014). Nodes on the original tree receiving less than 70% bootstrap support were collapsed. The final microsynteny tree was used to calculate distances covered by cluster distributions, as well as individual homolog groups.

### Sampling null models of gene cluster evolution

Empirically derived null models describing the background levels of “gene cluster” distributions were developed for each of the 12 largest taxonomic classes and each cluster size ranging from 4-24 genes (252 distributions in total). Briefly, for a given null distribution, a group of neighboring genes (i.e., query cluster) of size X was chosen at random from a randomly selected genome from taxonomic class Y. Hits to each gene in the query cluster were recovered from the proteome database using BLASTp. Genomes containing homologous clusters (i.e., containing hits to all genes in the query cluster, where no more than 6 intervening genes separated any hit from another) were then retrieved. To determine the phylogenetic distance associated with the query cluster distribution, the total branch length distance on the microsynteny tree connecting all genomes with homologous clusters was calculated and stored. The above sampling process was repeated 500 times for each null distribution.

### Locating Clusters through Unexpected Synteny

We have identified previously undetected gene clusters putatively involved in phenylpropanoid degradation using a method that posits that fungal genomes are spatially compartmentalized by function, similarly to those of bacteria (Snel et al. 2000; Rogozin et al. 2002; Rogozin et al. 2004; Szklarczyk et al. 2014; Zhao et al. 2014; Zaidi and Zhang 2016). Our method also draws upon the observation that microsynteny is often poorly conserved in fungal genomes (Hane et al. 2011), which has previously enabled biosynthetic MGC detection through the identification of conserved microsynteny among individuals of the same species (Takeda et al. 2014). Briefly, genes in regions surrounding “anchor genes” of interest were assigned to homolog groups. Clusters were then detected by comparing the phylogenetic distributions of homolog group combinations within these regions to distributions expected under null models of microsynteny evolution. More detailed steps are described below and in Supplementary Figure 1.

All homologs of an anchor gene query sequence of interest were retrieved using BLASTp. Twenty genes upstream and downstream of all anchor gene homologs (designated “anchor gene neighborhood”) were retrieved, compared against one another using UBLAST and sorted into homolog groups using OrthoMCL. Individual homolog groups whose distribution on the microsynteny species tree had a maximum pairwise distance below 0.95 were discarded. Within the set of all anchor gene neighborhoods, the set of unique combinations of 4 or more homolog groups that included the anchor gene (“cluster motifs”) was determined. The genomes in which each motif occurred were identified, with the condition that genes belonging to homolog groups in the motif never be separated by more than 6 intervening genes. For each genome in which a given motif occurred, the probability of observing the motif in that genome, given the size of the motif and the taxonomic class of the genome, was empirically estimated by determining the proportion of samples in the appropriate null distribution that cover a total distance on the microsynteny tree greater than or equal to the distance associated with the given motif, divided by the total number of samples in the null distribution. For example, to estimate the probability of observing a motif with 8 homolog groups in a Dothideomycete genome, we would compare the total distance associated with the observed motif to the null distribution of distances associated with size 8 clusters sampled from Dothideomycetes. For all such test, the test statistic is distance on the microsynteny tree, and the null hypothesis is that the phylogenetic distribution of a given cluster motif containing an anchor gene is consistent with background rates of microsynteny evolution. The null hypothesis is rejected for motifs with an estimated probability below the significance threshold of 0.05 in at least one genome, and all genes assigned to such motifs are designated clusters. Clusters with proteins that had fuNOG annotations or PFAM domains or GO term annotations associated with proteins known to exclusively participate in fungal secondary metabolite biosynthesis were excluded from further analysis (Supplementary Table 4). All annotations of clustered proteins were also manually inspected for evidence of exclusive participation in biosynthetic metabolism, and excluded if necessary.

### Cluster models and multi cluster model profiles (MCMPs)

Homology among clusters from the same cluster class (i.e., containing the same anchor gene) was determined by assessing similarities in homolog group content. Briefly, a matrix detailing the presence or absence of homolog groups in each cluster was used to calculate Bray-Curtis dissimilarity indices for all pairwise comparisons of clusters. Pairwise comparisons were then grouped using complete linkage clustering, and any clusters separated by under 0.6 distance units were assigned to the same cluster model. This cutoff was empirically determined after manual examination of the content of clusters assigned to the same model under various distance cutoffs. Homolog groups present in ≥75% of clusters assigned to a given model were then used to summarize that model. The above approach was also used to group fungal species with similar combinations of clusters into MCMPs by using a matrix detailing the presence or absence of homologous clusters (as determined by cluster model) among all species with two or more clusters. MCMPs observed in more than 5 species were used for enrichment analyses. All above analyses were performed using the vegan package in R (Oksanen et al. 2016).

### Network analyses

Amino acid sequences from all clustered homolog groups across the 13 cluster classes were combined into one set and then sorted into new homolog groups using OrthoMCL. Homolog groups that contained sequences from multiple cluster classes were designated as “shared”. Pairwise co-occurrences of all homolog groups in all clusters were determined, and visualized as a network with Cytoscape v.3.4.0 (Shannon et al. 2003). The network layout was determined solely by the AllegroLayout plugin with the Allegro Spring-Electric algorithm. Analyses of network modularity were performed with the spectral partitioning algorithm (Newman 2006), as implemented in MODULAR (Marquitti et al. 2014), and the probability of network modularity was estimated by randomly sampling the network 1000 times, twice.

### Enrichment tests

A one tailed Fisher's exact test, as implemented in the Text-NSP Perl module, was used to conduct all tests of enrichment (see Supplementary Figure 1 for examples of contingency table and formulae). Unless noted otherwise, gene cluster, cluster model or MCMP presence/absence was recorded at the species level, and counts of species were used to fill in contingency tables. Odds ratios were calculated to estimate the magnitude of the enrichment effect (Szumilas 2010). Contingency tables with a zero in least one cell had all cells incremented by 0.5 to avoid division by zero when calculating the odds ratio (Bradburn et al. 2007). In general, the odds ratio can be interpreted as the odds that fungi have a particular lifestyle, given they have a particular cluster-based feature (either cluster presence (Figure 1b), cluster model presence (Supplementary Figure 4) or MCMP assignment (Figure 2c). The precision of each odds ratio, except for those associated with contingency tables with a zero in at least one cell, was estimated by calculating the 95% confidence interval (Szumilas 2010). The general form of the null hypothesis is that particular features are not enriched in fungi with a particular ecological lifestyle. We rejected the null hypothesis at α = 0.05. As the Pezizomycotina possessed 89.4% of all clusters detected, only data from Pezizomycotina genomes were used in enrichments tests presented in Figure 1b, Supplementary Figure 4, and Figure 2c. Given the exploratory nature of this study, we did not correct for multiple testing in order to identify trends to follow up in later confirmatory studies.

## Code availability

Unless noted otherwise, all analyses were performed using a series of custom Perl scripts (Supplementary Figure 1). All scripts used to generate the presented results are available upon request. Fasta files of all recovered cluster models, as well as a script to recover cluster models identified in this analysis from a user-supplied genome of interest, are available for download at https://github.com/egluckthaler/cluster_retrieve

## Acknowledgements and funding

This work was supported by funds from The Ohio Agricultural Research and Development Center at The Ohio State University (EGT, JCS), The National Science Foundation (DEB-1638999, JCS) and the Fonds de recherche du Québec-Nature et technologies (EGT). All computational work was conducted on the Ohio State Supercomputer. We thank [complete author list to be determined] for providing access to unpublished genome data produced by the U.S. Department of Energy Joint Genome Institute, a DOE Office of Science User Facility, supported by the Office of Science of the U.S. Department of Energy under Contract No. DE-AC02-05CH11231.

## Author contributions

EGT and JCS conceptualized the work and wrote the manuscript. EGT developed and performed all analyses.

## Competing financial interests

The authors have declared that no competing interests exist.

